# Melittin intervention induces lncRNA response, mitochondrial dysfunction, and cell proliferation in murine cervical cancer cells

**DOI:** 10.64898/2026.08.18.745438

**Authors:** Ronghua Zhang, Haiwen Zhuo, Yunzhen Yang, Kaiyao Zhang, Mengyi Wang, Jianrong Jiang, Yueliang Li, Jianfeng Qiu, Dafu Chen, Tizhen Yan, Rui Guo

## Abstract

Melittin exhibits antitumor activity in cervical cancer models, yet the long non-coding RNA (lncRNA) response and associated regulatory networks remain poorly understood. Here, strand-specific RNA-seq data from melittin-treated and untreated U14 murine cervical cancer cells were analyzed to characterize melittin-responsive lncRNAs and explore their potential functional associations. A total of 28,162 lncRNAs were identified, including 27,307 known and 855 novel transcripts. Differential expression analysis revealed 404 differentially expressed lncRNAs (DElncRNAs), comprising 191 upregulated and 213 downregulated lncRNAs, most of which were predicted to localize to the cytoplasm or nucleus. *Cis*-target analysis identified 52 neighboring mRNAs as putative targets of 46 DElncRNAs. Functional enrichment highlighted mitochondrial electron transfer and redox-related processes, including the mitochondrial electron transfer flavoprotein complex, electron-transferring-flavoprotein dehydrogenase activity, ubiquinone binding, and quinone binding. In parallel, melittin induced mitochondrial membrane depolarization and increased intracellular reactive oxygen species accumulation in U14 cells. Co-expression analysis further identified 138 lncRNAs coexpressed with 161 mRNAs, which were enriched in chromatin remodeling, DNA replication, and DNA repair. EdU incorporation decreased with increasing melittin concentrations, indicating suppression of DNA synthesis and proliferative activity. RT-qPCR analysis confirmed the expression trends of selected DElncRNAs. Collectively, these findings demonstrate extensive remodeling of the lncRNA landscape in melittin-treated U14 cells and suggest that melittin-responsive lncRNA-mRNA networks are associated with mitochondrial redox disruption and impaired DNA synthesis. This study provides a transcriptomic framework for identifying candidate lncRNA-mRNA regulatory axes underlying the antitumor response to melittin.

## 1. Introduction

Cervical cancer remains a major gynecological malignancy worldwide. Although vaccination, screening, surgery, radiotherapy, chemotherapy, targeted therapy, and immunotherapy have substantially improved its prevention and clinical management, recurrent, metastatic, and treatment-resistant disease remains difficult to control. These clinical challenges highlight the need to identify novel therapeutic agents with improved efficacy and acceptable safety profiles. In this context, natural bioactive compounds have attracted increasing attention because of their structural diversity, multitarget biological activities, and capacity to suppress tumor cell proliferation, induce cell death, and modulate treatment-related responses.

Melittin, a major peptide component of bee venom, has emerged as a potential anticancer molecule because of its ability to inhibit tumor cell growth and promote cancer cell death[1] [1]. Previous studies have demonstrated that melittin suppresses the proliferation, migration, and invasion of various cancer cell types and induces cell death [2] through mechanisms involving mitochondrial dysfunction, oxidative stress, and apoptosis-related signaling [3][2,3]. For example, melittin can directly interact with phospholipids in the inner mitochondrial membrane, disrupting membrane lipid organization and subsequently impairing mitochondrial membrane potential and bioenergetic function[4] [4]. Similarly, melittin has been shown to activate Bax/Bcl-2-associated mitochondrial apoptotic signaling in A549 lung cancer cells [5][5]. Although accumulating evidence supports the antitumor activity of melittin, including its effects in human cervical cancer cell lines such as HeLa and C33A, previous studies have predominantly focused on cellular phenotypes and protein-associated signaling mechanisms. The transcriptomic regulatory response, particularly that mediated by long non-coding RNAs (lncRNAs), to direct melittin treatment in cervical cancer cells remains poorly understood.

Long non-coding RNAs are transcripts longer than 200 nucleotides with limited protein-coding potential and function as important regulators of gene expression at the epigenetic, transcriptional, and post-transcriptional levels. Accumulating evidence indicates that lncRNAs participate in multiple cancer-related processes, including sustained proliferation, evasion of cell death, invasion and metastasis, tumor–microenvironment interactions, and therapeutic resistance[6, 7] [6,7]. In cervical cancer, aberrantly expressed lncRNAs have been implicated in the regulation of tumor cell proliferation, apoptosis, metastatic behavior, and treatment response. For example, LINC00460 promotes cervical cancer cell proliferation and inhibits apoptosis through the miR-503-5p axis[8], whereas LINC00657 promotes malignant progression by regulating the miR-30s/SKP2 axis[9]. In addition, NCK1-AS1 has been associated with cisplatin resistance through regulation of the miR-134-5p/MSH2 axis[10]. These findings highlight the functional importance of lncRNA-mediated regulatory networks in cervical cancer. With the development of high-throughput RNA sequencing, systematic lncRNA profiling has also become an effective approach for identifying treatment-responsive regulatory molecules and their associated signaling networks[11-13]. Transcriptomic analyses have, for example, been used to characterize lncRNA-associated responses to nanocurcumin treatment in colorectal cancer cells and to identify competing endogenous RNA networks associated with endocrine resistance in breast cancer cells[14-16]. However, the lncRNA regulatory landscape associated with the response of cervical cancer cells to melittin remains largely unexplored.

In our previous study, melittin-treated and untreated murine cervical carcinoma U14 cells were subjected to strand-specific cDNA library construction and high-throughput RNA sequencing, generating high-quality transcriptomic datasets for comparative investigation of melittin-induced transcriptional alterations[17]. Building on these datasets, the present study systematically identified and characterized lncRNAs, evaluated their structural features and expression profiles, and investigated differentially expressed lncRNAs (DElncRNAs) and their potential regulatory functions. To complement the transcriptomic analyses with phenotypic and functional evidence, we further assessed cell proliferation by EdU incorporation, intracellular reactive oxygen species (ROS) accumulation, mitochondrial membrane potential, and the expression of selected transcripts by RT-qPCR. Collectively, this study aims to characterize the lncRNA-associated molecular response of U14 cervical cancer cells to melittin and to provide a foundation for further elucidating the lncRNA-mediated regulatory mechanisms underlying its antitumor effects.

## 2. Materials and Methods

### 2.1 Cell culture and melittin treatment

The murine cervical carcinoma U14 cell line was purchased from Immocell Biotechnology (Xiamen, China) and cultured in high-glucose Dulbecco’s Modified Eagle Medium (DMEM; Gibco, USA) supplemented with 10% fetal bovine serum (FBS; Gibco, USA) and 1% penicillin–streptomycin (Yeasen, Shanghai, China). Cells were maintained at 37°C in a humidified atmosphere containing 5% CO_2_, and used for subsequent experiments during the logarithmic growth phase. Melittin (purity >99%) was purchased from Selleck Chemicals (Houston, TX, USA). Lyophilized melittin was dissolved in sterile water to prepare a 4 mg/mL stock solution, aliquoted, and stored at −80°C until use. Before each experiment, the stock solution was diluted with complete DMEM to the indicated working concentrations.

### 2.2 EdU incorporation assay

U14 cells were seeded in 24-well plates and allowed to adhere overnight. The cells were subsequently exposed to melittin at 0, 2, 4, 6, or 8 μg/mL for 20 min. DNA synthesis was assessed using a Cell-Light™ EdU Apollo594 In Vitro Imaging Kit (RiboBio, Guangzhou, China). Following a 20 min incubation with EdU labeling, the cells were processed according to the manufacturer’s protocol, and nuclei were counterstained with Hoechst 33342 (Yeasen, Shanghai, China). Fluorescence images were acquired using an inverted fluorescence microscope (ECLIPSE Ts2; Nikon, Tokyo, Japan). EdU incorporation was quantified as the percentage of EdU-positive nuclei relative to the total number of Hoechst-positive nuclei. Three independent experiments were performed.

### 2.3 ROS measurement

U14 cells were treated with melittin at 0, 2, 4, 6, or 8 μg/mL for 20 min. Intracellular ROS levels were subsequently evaluated using the fluorescent probe 2′,7′-dichlorodihydrofluorescein diacetate (DCFH-DA; Beyotime Biotechnology, Shanghai, China) according to the manufacturer’s protocol. Fluorescence images were acquired using the fluorescence microscope described above. The fluorescence intensity of 2′,7′-dichlorofluorescein (DCF) was quantified using ImageJ software (v1.53; National Institutes of Health, Bethesda, MD, USA)[18]. Three independent experiments were conducted.

### 2.4 Detection of mitochondrial membrane potential

U14 cells were exposed to melittin at 0, 2, 4, 6, or 8 μg/mL for 20 min. Carbonyl cyanide 3-chlorophenylhydrazone (CCCP)-treated cells were included as a positive control. Mitochondrial membrane potential (ΔΨm) was evaluated using a JC-1 Mitochondrial Membrane Potential Assay Kit (Beyotime Biotechnology, Shanghai, China). Following JC-1 staining, fluorescence images were obtained using the fluorescence microscope described above. Red and green fluorescence intensities were quantified using ImageJ (v1.53), and ΔΨm was expressed as the ratio of red-to-green fluorescence intensity. Measurements were obtained from three independent experiments.

### 2.5 RNA-seq data source

Strand-specific RNA-sequencing datasets generated from U14 cells treated with 4 μg/mL melittin (T) and untreated control cells receiving an equal volume of complete culture medium (C), with three biological replicates per group, were obtained from our previous study. RNA extraction, ribosomal RNA depletion, strand-specific library construction, paired-end 150-bp sequencing, and quality control of the sequencing data were performed as previously described [17]. The high-quality clean reads generated from these experiments were used for subsequent transcriptome reconstruction and lncRNA analyses in the present study.

### 2.6 Transcript assembly and expression quantification

Clean reads were aligned to the *Mus musculus* reference genome GRCm39 using HISAT2 (v2.2.1). The corresponding gene annotation file was obtained from Ensembl release 112. Transcript assembly and reconstruction were performed using StringTie (v2.1.6) based on the resulting alignments. Individual transcript assemblies were subsequently integrated to generate a unified transcript annotation for downstream analyses.

Transcript expression was quantified against the final transcript annotation. Fragments per kilobase of transcript per million mapped reads (FPKM) were used to characterize transcript abundance and assess expression profiles and sample correlations, whereas raw read counts were used for differential expression analysis.

### 2.7 Identification of lncRNAs

Transcripts annotated as known protein-coding mRNAs and transcripts shorter than 200 nucleotides were excluded from the reconstructed transcript set. The coding potential of the remaining unannotated transcripts was independently evaluated using the Coding Potential Calculator (CPC, v0.9-r2) and Coding-Non-Coding Index (CNCI, v2.0). Transcripts predicted to possess coding potential were excluded, whereas transcripts consistently classified as non-coding by both algorithms were retained as novel lncRNA candidates. These novel lncRNAs were combined with reference-annotated lncRNAs for subsequent expression profiling and differential expression analyses.

### 2.8 Screening of DElncRNAs

Differential expression analysis between melittin-treated and untreated U14 cells was performed using raw read counts with DESeq2 (v1.22.2). *P*-values were adjusted for multiple comparisons using the Benjamini–Hochberg procedure. lncRNAs with an absolute log_2_ fold change ≥ 1 and an adjusted *P* value < 0.05 were defined as differentially expressed lncRNAs (DElncRNAs).

### 2.9 Subcellular localization prediction

The subcellular localization of DElncRNAs was predicted using RNAlight (v1.0), iLoc-LncRNA (v2.0), and DeepLncLoc. RNAlight predicts nuclear or cytoplasmic localization using a LightGBM-based classification model, whereas iLoc-LncRNA employs a support vector machine-based approach. DeepLncLoc predicts localization probabilities across multiple subcellular compartments using a deep learning-based framework. For DeepLncLoc, prediction confidence was further evaluated according to the difference between the highest and second-highest localization scores. Concordant localization assignments obtained from the three prediction tools were considered high-confidence predictions for subsequent analyses.

### 2.10 Prediction of *cis*-regulated target genes of DElncRNAs

Potential *cis*-regulated target genes of DElncRNAs were identified according to their genomic proximity to protein-coding genes. Protein-coding genes located within 100 kb upstream or downstream of a DElncRNA were considered candidate *cis*-target genes. Differentially expressed mRNAs (DEmRNAs) within these genomic regions were subsequently retained to construct DElncRNA–mRNA regulatory pairs. The resulting *cis*-regulatory network was visualized using Cytoscape (v3.10.3). Gene Ontology (GO) and Kyoto Encyclopedia of Genes and Genomes (KEGG) pathway enrichment analyses were subsequently performed for the identified target genes.

### 2.11 Identification of Co-Expressed mRNAs Associated with DElncRNAs

Potential co-expression relationships between DElncRNAs and mRNAs were evaluated by Pearson correlation analysis using their expression profiles across melittin-treated and untreated U14 samples. DElncRNA–mRNA pairs meeting the predefined correlation criteria were retained and used to construct a co-expression network, which was visualized using Cytoscape (v3.10.3). GO and KEGG enrichment analyses were subsequently performed for the co-expressed mRNAs.

### 2.12 RT-qPCR validation

To evaluate the reliability of the RNA-seq results, five DElncRNAs, including ENSMUST00000243162, MSTRG.9633.2, MSTRG.12176.6, MSTRG.7888.6, and MSTRG.6027.5, were selected for RT-qPCR validation. Specific primers were designed using Primer Premier 6.0 and synthesized by Sangon Biotech (Shanghai, China). Primer sequences are provided in Supplementary Table S1.

Total RNA was extracted from U14 cells and reverse-transcribed into complementary DNA (cDNA) according to the manufacturers’ protocols. RT-qPCR was performed using SYBR Green chemistry in a final reaction volume of 20 μL. The amplification conditions consisted of an initial denaturation at 95°C for 1 min, followed by 40 cycles of 95°C for 10 s and 55°C for 30 s. *GAPDH* (GenBank accession no. NM_008084.4) was used as the internal reference gene. Each sample was analyzed in technical triplicate, and relative lncRNA expression was calculated using the 2^−ΔΔCt^ method[19] [22].

### 2.13 Statistical analysis

Data from the EdU incorporation, intracellular ROS, and mitochondrial membrane potential assays are presented as the mean ± standard deviation (SD) from three independent experiments. Differences among the five melittin concentration groups were analyzed using one-way analysis of variance (ANOVA), followed by Dunnett’s multiple-comparisons test using the untreated group as the reference. RT-qPCR data between melittin-treated and untreated groups were compared using an unpaired two-tailed Student’s *t*-test. Statistical analyses were performed using GraphPad Prism 10.0 (GraphPad Software, Boston, MA, USA). A *P* value < 0.05 was considered statistically significant.

## 3. Results

### 3.1 Global identification of lncRNAs in U14 murine cervical cancer cells

Based on sequencing data from all melittin-treated and control U14 cell samples, CPC and CNCI predicted 28,296 and 28,281 lncRNAs, respectively, of which 28,162 were commonly identified by both methods (Fig. 1a). Among the redundant lncRNAs, 27,307 (96.96%) were annotated as known lncRNAs, while 855 (3.04%) were classified as novel one (Fig. 1b). Among the 855 novel lncRNAs, intergenic lncRNAs were the most abundant, accounting for 519 transcripts (60.70%), followed by intronic lncRNAs (212, 24.80%). Novel isoforms, exonic-overlapping lncRNAs, and antisense lncRNAs accounted for 60 (7.02%), 44 (5.15%), and 20 (2.34%) transcripts, respectively (Fig. 1c).

**Fig 1.**
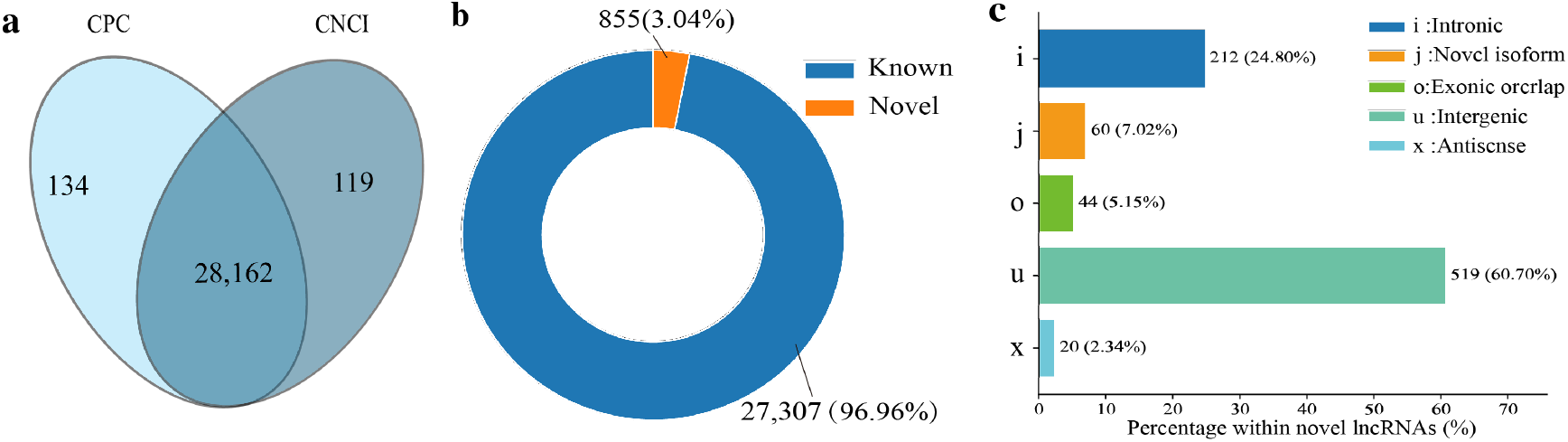
Identification and classification of lncRNAs in melittin- and un-treated murine cervical cancer U14 cells. (**a**) Intersection of candidate lncRNAs predicted by CPC and CNCI; (**b**) ratios of known and novel lncRNAs; (**c**) distribution of various categories of novel lncRNAs

### 3.2 Subcellular Localization Prediction of DElncRNAs

In total, 404 DElncRNAs were identified in U14 cells following melittin intervention, including 381 known and 23 novel lncRNAs. Among these, there were 191 up- and 213 down-regulated lncRNAs (Fig. 2a, see also Table S2). Subcellular localization prediction showed that majority of DElncRNAs were distributed in the cytoplasm. Specifically, by using RNAlight, 203 down- and 187 up-regulated lncRNAs were predicted to localize in cytoplasm, whereas 10 down- and 4 up-regulated ones were assigned to exosomes (Fig. 2b). Consistently, based on iLoc-LncRNA, cytoplasm was predicted as the major localization site followed by the nucleus, a small proportion of lncRNAs were predicted to distributed in ribosomes and exosomes (Fig. 2c). As shown in Fig. 2d, the localization probability heatmap further confirmed that DElncRNAs were mainly distributed in cytoplasm and nucleus. In addition, Venn analysis further revealed that 9 exosome-localized lncRNAs were commonly predicted by iLoc-LncRNA and DeepLncLoc, whereas 261 cytoplasm-localized lncRNAs were shared between RNAlight and iLoc-LncRNA. For nuclear localization, one lncRNA was concordantly predicted by all three algorithms, with additional pairwise overlaps observed among the prediction tools (Fig. 2e–g).

**Fig 2.**
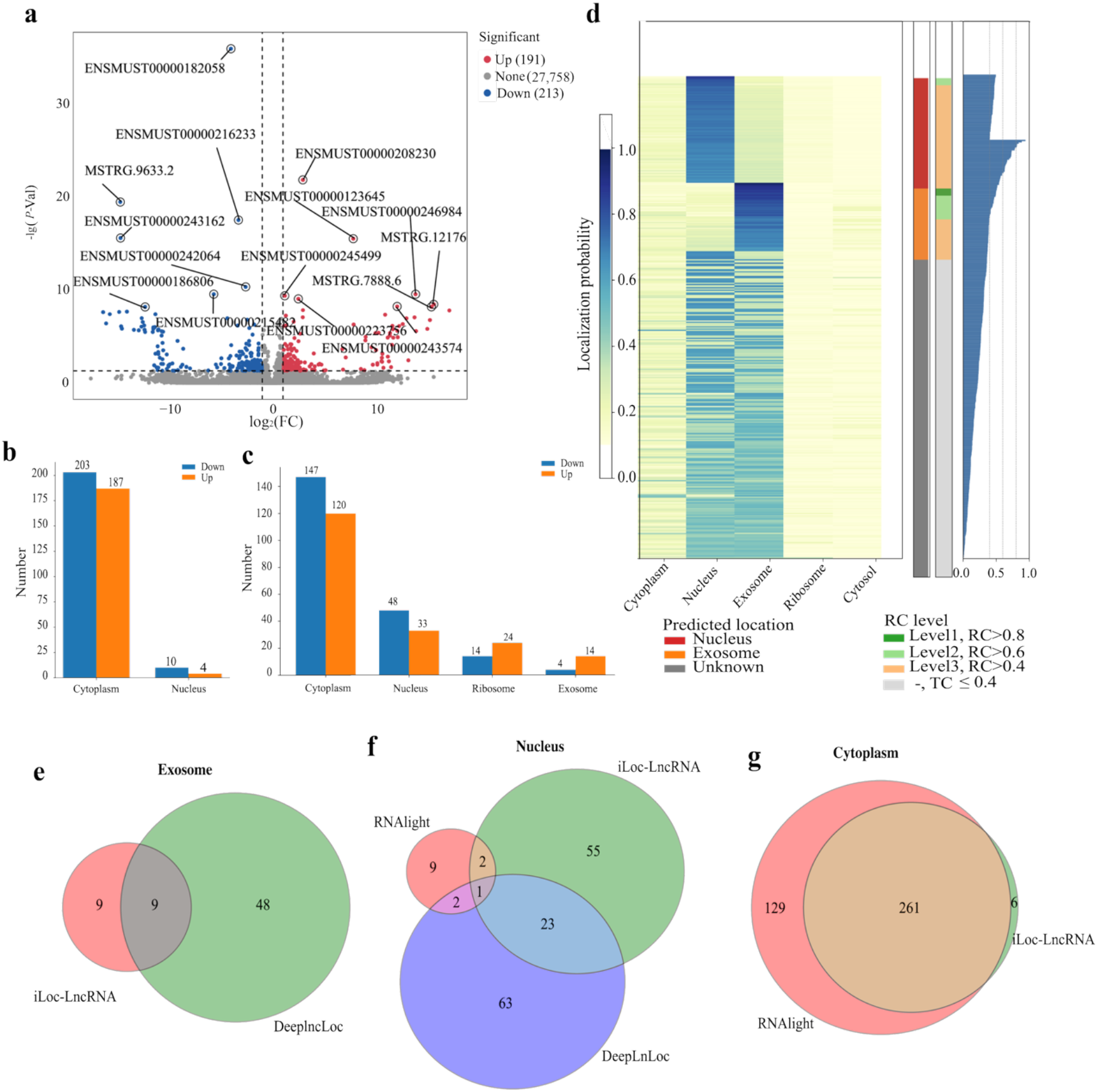
Subcellular localization prediction of differentially expressed lncRNAs. (a) Volcano plot of differentially expressed lncRNAs (DElncRNAs) between the melittin treated and untreated groups; (b-c) subcellular localization of DElncRNAs predicted by RNAlight and iLoc-LncRNA; (d) heatmap of localization probabilities and reliability scores across subcellular compartments; (e-g) Venn diagram showing the overlap of exosome-, nucleus-, and cytoplasm-localized lncRNAs.

### *3*.*3 Cis*-acting role of DElncRNAs in U14 murine cervical cancer cells following melittin treatment

A total of 46 DElncRNAs were predicted to cis-regulate 52 mRNAs (Fig. 3a), which were enriched in 334 biological process-, 221 cellular component-, and 192 molecular -related GO terms, such as lipid tube assembly involved in organelle fusion, negative regulation of mRNA modification, mitochondrial electron transfer flavoprotein complex, mRNA editing complex, electron-transferring-flavoprotein dehydrogenase activity, and ubiquinone binding (Fig. 3b). Notably, a subseries of functional terms relative to mitochondrial electron transport and redox metabolism were enriched by target mRNAs, including the mitochondrial electron transfer flavoprotein complex, electron-transferring-flavoprotein dehydrogenase activity, ubiquinone binding, and quinone binding. KEGG enrichment analysis revealed four significantly enriched pathways, such as RNA polymerase, nucleotide excision repair, cytosolic DNA-sensing pathway, and ribosome biogenesis in eukaryotes (Fig. 3c).

**Fig 3.**
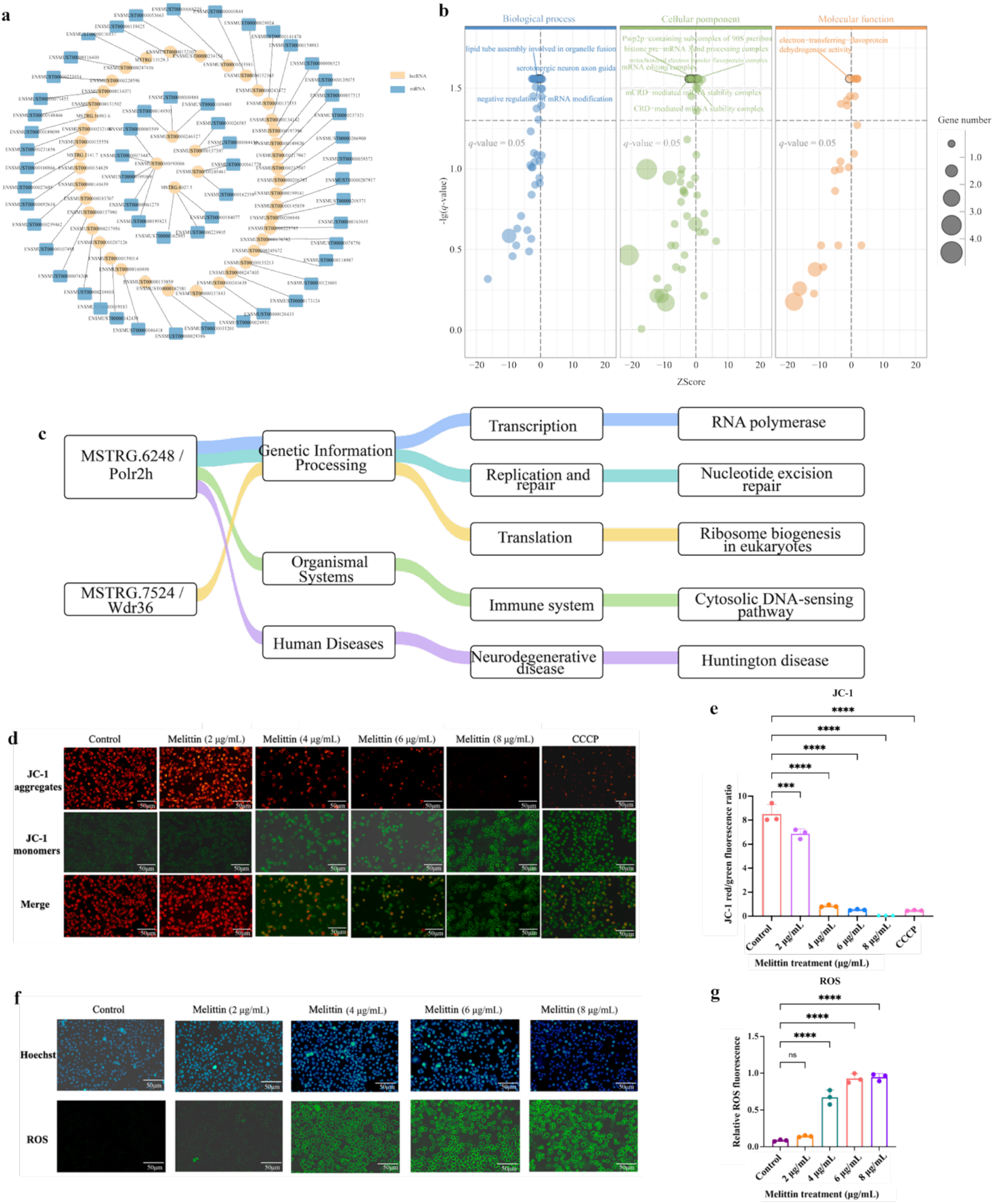
Functional enrichment of cis-targeted mRNAs and melittin-induced mitochondrial depolarization in U14 cells. (a) Interaction network between DElncRNAs and their *cis-*acting mRNAs; (b) bubble diagrams of GO terms enriched by *cis*-acting mRNAs; (c) Sankey diagram showing KEGG pathway classification and enrichment of *cis*-acting mRNAs; (d) representative JC-1 fluorescence images of U14 cells treated with 0, 2, 4, 6 or 8 μg/mL melittin for 20 min, with CCCP as a positive control; (e) quantification of the JC-1 red-to-green fluorescence intensity ratio; (f) representative fluorescence images of intracellular ROS after melittin treatment; (g) quantification of intracellular ROS fluorescence intensity.

Additionally, JC-1 staining showed that melittin progressively reduced red aggregate fluorescence while increasing green monomer fluorescence, indicating the melittin-caused loss of mitochondrial membrane potential (Fig. 3d). Likewise, the JC-1 red/green fluorescence ratio was significantly decreased at all tested concentrations, with a marked reduction at 4–8 μg/mL and values at 6–8 μg/mL approaching those of the CCCP positive control (Fig. 3e).

Moreover, DCFH-DA fluorescence increased markedly in cells treated with 4, 6 and 8 μg/mL melittin, suggestive of enhanced intracellular ROS accumulation (Fig. 3f-g).

### 3.4 Investigation of co-expressed mRNAs of DElncRNAs and inhibition of DNA synthesis by melittin

Here, 138 lncRNAs were detected to be co-expressed with 161 mRNAs, and representative lncRNA–mRNA interactions were visualized as a co-expression network (Fig.4a). These co-expressed mRNAs were involved in 302 functional terms like transcriptional regulation, chromatin organization, and DNA binding. Several GO terms were associated with DNA replication and genome stability, such as chromatin remodeling, regulation of DNA replication and DNA repair (Fig. 4b). In addition, the co-expressed targets were engaged in an array of KEGG pathways relevant to herpes simplex virus 1 infection, alcoholism, and systemic lupus erythematosus showed the highest statistical significance (Fig. 4c). Further examination revealed that several enriched pathways were associated with cellular stress and cell proliferation, including neutrophil extracellular trap formation, viral carcinogenesis and ATP-dependent chromatin remodeling. To determine whether these functional alterations were accompanied by changes in inhibition of DNA synthesis by melittin and cell proliferation, EdU incorporation was examined in U14 cells following treatment with different concentrations of melittin, the results suggested that EdU fluorescence and the proportion of EdU-positive cells progressively decreased with increasing melittin concentrations (Fig. 4d). Compared with the control group, 2 μg/mL melittin caused no significant alteration, whereas treatment with 4, 6 and 8 μg/mL significantly reduced EdU incorporation, with the most pronounced inhibition observed at 6 and 8 μg/mL (Fig. 4e).

**Fig 4.**
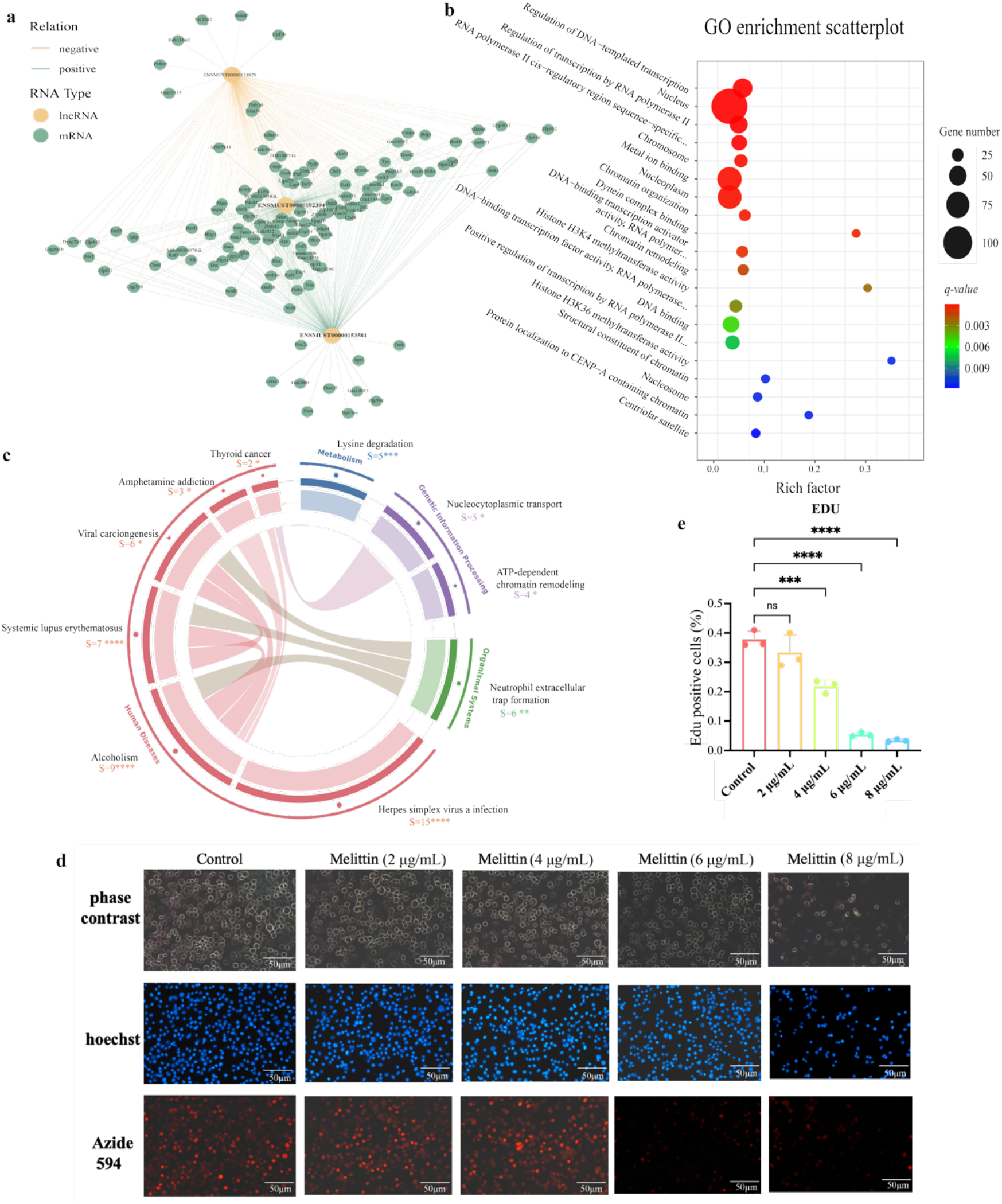
Investigation of co-expressed mRNAs of DElncRNAs and validation of melittin-induced inhibition of DNA synthesis in U14 cells. (a) Representative co-expression network of DElncRNAs and the co-expressed mRNA; (b) bubble plot of GO terms enriched by the co-expressed mRNAs; (c) Chord diagram showing the relationships between representative significantly enriched KEGG pathways and their corresponding co-expressed mRNAs; (d) Representative EdU staining images of U14 cells treated with 0–8 μg/mL melittin for 20 min. (e) Quantification of EdU-positive cells. Data are presented as mean ± SD (n = 3). ns, not significant, \*\*\**P* < 0.001, \*\*\*\**P* < 0.0001.

### 3.5 RT-qPCR validation of DElncRNA expression

Five DElncRNAs were randomly selected for RT-qPCR validation, the results demonstrated that the expression trends of these lncRNAs were consistent with the RNA-seq data, which verified the reliability of the used transcriptome data.

## Discussion

As important regulators of gene expression, lncRNAs have been widely implicated in cancer progression and treatment response [6, 20]. Previous lncRNA profiling studies in cervical cancer have commonly inferred the potential functions of dysregulated lncRNAs through their associated protein-coding genes, together with GO and KEGG enrichment analyses and lncRNA – mRNA network construction[21]. In the present study, 404 DElncRNAs were identified in U14 cells following melittin intervention (Fig. 2a), indicating a marked lncRNA transcriptional response to melittin. Subcellular localization prediction further suggested that these DElncRNAs were predominantly localized to the cytoplasm, with a subset predicted to occur in the nucleus (Fig. 2b–d). This distribution is of potential functional relevance because cytoplasmic lncRNAs commonly participate in the regulation of mRNA stability, translation, and signaling, whereas nuclear lncRNAs are more frequently associated with chromatin organization and transcriptional regulation[22]. Moreover, the concordant expression trends obtained by RNA-seq and RT-qPCR for the selected DElncRNAs supported the reproducibility of the differential expression profiles (Fig. 5).

**Fig 5.**
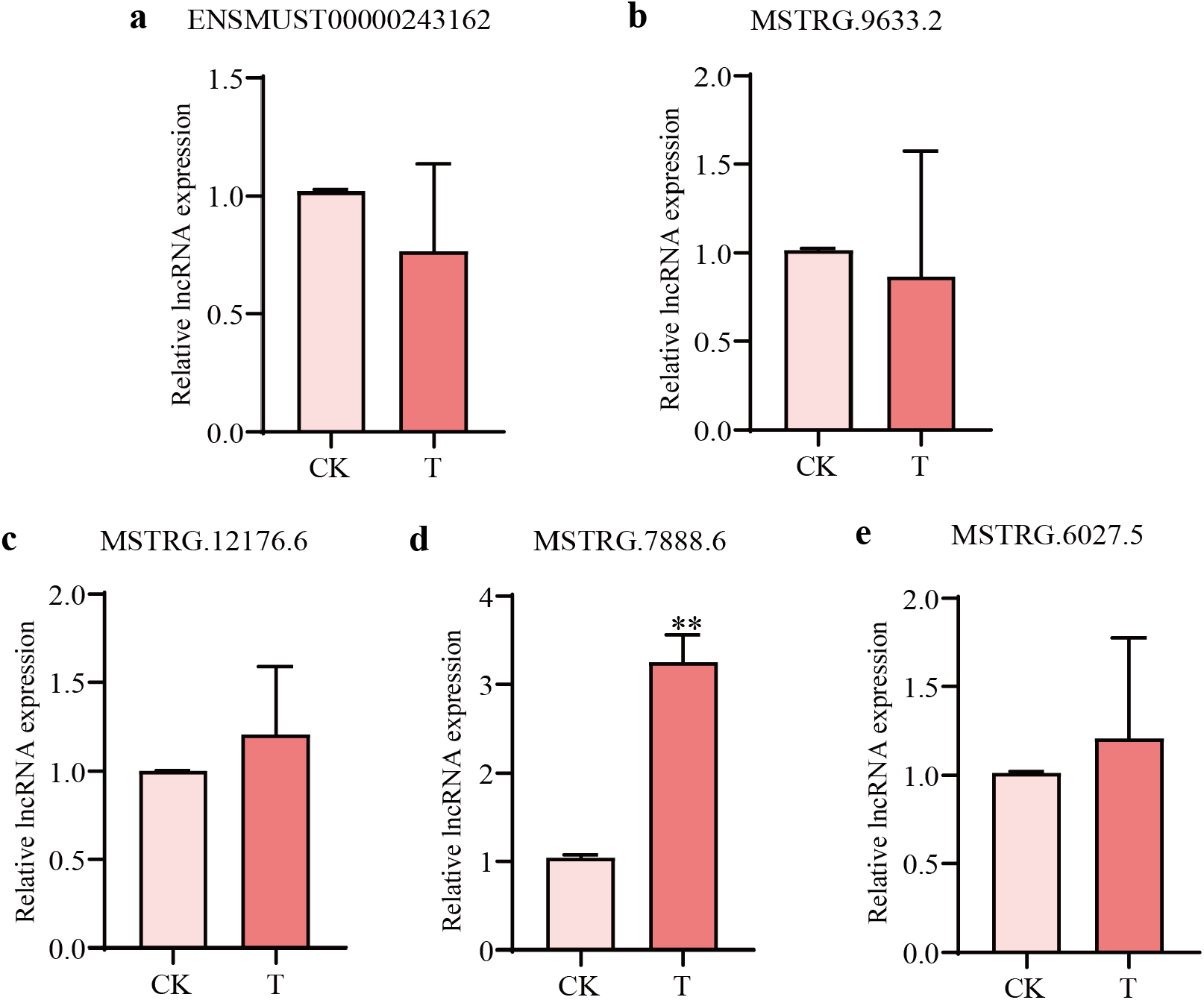
RT-qPCR verification of five DElncRNAs. Data are presented as Mean ± SD and subjected to Student’s *t* test (ns: *P*>0.05, *: *P*<0.05, **: *P*<0.01, ***: *P*<0.001, ****: *P*<0.0001).

The *cis*-target analysis provided further insight into the potential biological functions associated with melittin-responsive DElncRNAs. A total of 46 DElncRNAs were predicted to *cis*-regulate 52 neighboring mRNAs, forming a putative DElncRNA–mRNA regulatory network (Fig. 3a). Functional enrichment analysis identified several terms related to mitochondrial electron transfer and redox processes, including the mitochondrial electron transfer flavoprotein complex, electron-transferring-flavoprotein dehydrogenase activity, ubiquinone binding, and quinone binding (Fig. 3b). These functions converge on flavoprotein-dependent electron transfer and electron delivery to the ubiquinone pool, both of which are closely associated with mitochondrial redox homeostasis. In addition, KEGG enrichment identified pathways related to RNA polymerase, nucleotide excision repair, cytosolic DNA sensing, and ribosome biogenesis (Fig. 3c), suggesting that the predicted *cis*-associated genes may participate in broader transcriptional and cellular stress responses. Collectively, these bioinformatic results indicate a potential association between melittin-responsive DElncRNAs and mitochondrial/redox-related processes, although the predicted regulatory relationships require direct experimental validation.

In parallel with these transcriptomic signatures, melittin intervention resulted in a progressive decrease in mitochondrial membrane potential, together with a marked increase in intracellular ROS levels at 4–8 μg/mL (Fig. 3d–g). Because these changes were detected after only 20 min of melittin exposure, mitochondrial depolarization and ROS accumulation may represent relatively early cellular responses to melittin rather than solely secondary consequences of prolonged growth inhibition. Previous studies have shown that lncRNAs regulate mitochondrial bioenergetics, dynamics, and stress adaptation by regulating mitochondrial or nuclear-encoded factors[23]. For example, HOXA11os regulates mitochondrial function and cellular metabolism, whereas Caren and CARL have been implicated in mitochondrial bioenergetics and dynamics through distinct lncRNA-dependent mechanisms[24]. Melittin has also been reported to directly disrupt mitochondrial function, reduce mitochondrial membrane potential and promote ROS generation in several cancer models[5, 25]. Therefore, the convergence of mitochondrial- and redox-related *cis*-target enrichment with the JC-1 and ROS phenotypes observed in the present study supports an association between melittin-responsive lncRNAs and mitochondrial redox disturbance in U14 cells. However, the current evidence remains associative, and whether individual DElncRNAs directly participate in mitochondrial depolarization or ROS accumulation remains to be determined.

In addition to the mitochondrial and redox-associated signatures, mRNAs co-expressed with DElncRNAs were enriched in biological processes related to chromatin remodeling, regulation of DNA replication, and DNA repair (Fig. 4a–c). These processes are closely interconnected with genome maintenance and cell-cycle progression, as appropriate chromatin organization is required for coordinated DNA replication and repair[26]. Correspondingly, EdU incorporation progressively decreased with increasing melittin concentrations, with significant reductions observed following treatment with 4, 6, and 8 μg/mL melittin (Fig. 4d, e). These observations demonstrate that melittin suppresses DNA synthesis in U14 cells and are consistent with the enrichment of DNA replication- and chromatin-associated functions in the DElncRNA co-expression analysis. Previous studies have also implicated lncRNAs in the regulation of proliferative activity in cervical cancer. For example, CCHE1 has been reported to promote cervical cancer cell proliferation through regulation of PCNA, whereas PVT1 contributes to the proliferative phenotype of cervical cancer cells[27, 28]. In other cancer models, melittin-responsive lncRNAs have likewise been associated with alterations in tumor cell proliferation. Taken together, these observations raise the possibility that melittin-responsive lncRNA– mRNA networks are associated with reduced DNA synthesis and proliferative activity in U14 cells. Nevertheless, perturbation of individual DElncRNAs will be required to determine whether these transcripts directly contribute to the antiproliferative response to melittin.

Collectively, the present study reveals a broad lncRNA response of U14 cells to melittin intervention and identifies two major functional features associated with this response. Predicted *cis*-associated mRNAs were linked to mitochondrial electron transfer and redox-related processes, whereas DElncRNA-co-expressed mRNAs were enriched in chromatin remodeling, DNA replication, and DNA repair. These transcriptomic signatures were accompanied by mitochondrial membrane depolarization, ROS accumulation, and reduced DNA synthesis, providing complementary molecular and cellular evidence for the response of U14 cells to melittin. Importantly, the present analyses identify candidate lncRNA–mRNA associations rather than establishing direct regulatory relationships. Future studies incorporating gain- and loss-of-function manipulation of selected DElncRNAs, together with assessment of their predicted target genes and corresponding mitochondrial, redox, and proliferative phenotypes, will be required to define the specific regulatory axes involved in the antitumor effects of melittin.

## Author Contributions

Conceptualization, R.Z., H.Z., Y.Y., D.C., T.Y. and R.G.; methodology, R.Z., H.Z., Y.Y. and K.Z.; software, Y.Y., K.Z. and M.W.; validation, R.Z., H.Z. and Y.Y.; formal analysis, H.Z. and M.W.; investigation, K.Z., M.W., J.J. and Y.L.; resources, J.Q., D.C., T.Y. and R.G.; data curation, R.Z., H.Z., Y.Y. and K.Z.; writing—original draft preparation, R.Z., H.Z. and Y.Y.; writing—review and editing, J.Q., D.C., T.Y. and R.G.; visualization, Y.Y. and M.W.; supervision, D.C., T.Y. and R.G.; project administration, J.Q., D.C., T.Y. and R.G.; funding acquisition, D.C., T.Y. and R.G. All authors have read and agreed to the published version of the manuscript.

## Institutional Review Board Statement

Not applicable.

## Informed Consent Statement

Not applicable.

## Data Availability Statement

The RNA-sequencing datasets analyzed in this study are publicly available in the Genome Sequence Archive (GSA) of the National Genomics Data Center (NGDC) under BioProject accession PRJCA068439. The datasets were previously reported in our bioRxiv preprint (doi: 10.64898/2026.07.25.739748, posted on 28 July 2026) and are available under a CC BY 4.0 International license.

## Acknowledgments

During the preparation of this manuscript, the authors used ChatGPT (OpenAI) for language editing and improvement of scientific writing. The authors reviewed and edited the output and take full responsibility for the content of this publication.

## Abbreviations

The following abbreviations are used in this manuscript:

ANOVA: Analysis of variance
CCCP: Carbonyl cyanide 3-chlorophenylhydrazone
CDNA: Complementary DNA
CNCI: Coding-Non-Coding Index
CPC: Coding Potential Calculator
DCF: 2’7’-Dichlorofluorescein
DCFH-DA: 2’7’-Dichlorodihydrofluorescein diacetate
DEInCRNA: Differentially expressed long non-coding RNA
DMEM: Dulbecco’s Modified Eagle Medium
EdU: 5-Ethynyl-2’-deoxyuridine
FBS: Fetal bovine serum
FPKM: Fragments per kilobase of transcript per million mapped reads
GO: Gene Ontology
GTF: Gene Transfer Format
JC-1: 5,5’6,6’-Tetrachloro-1,1’3,3’tetraethylbenzimidazolylcarbocyanine iodide
KEGG: Kyoto Encyclopedia of Genes and Genomes
IncRNA: Long non-coding RNA
mRNA: Messenger RNA
RC: Reliability coefficient
RNA-seq: RNA sequencing
ROS: Reactive oxygen species
RT-qPCR: Reverse transcription quantitative polymerase chain reaction
SD: Standard deviation
AΨm: Mitochondrial membrane potential

