## Supplemental table 1 for "Melittin intervention induces lncRNA response, mitochondrial dysfunction, and cell proliferation in murine cervical cancer cells"

**Table S1.** Sequences of primers qPCR detection.

| **Gene** | **Sequence (5′-3′)** |
| --- | --- |
| *Gadph* | F: AGGTCGGTGTGAACGGATTTG |
|  | R: GGGGTCGTTGATGGCAACA |
| ENSMUST00000243162 | F: ATGGACTGTGAGCCTGTGT |
|  | R: TGTGTAAGCCAGAGAACAACA |
| MSTRG.9633.2 | F: TCCAAGTGACGAGTGACATAG |
|  | R: TCTTCTGGTGAGGACATTCTG |
| MSTRG.12176.6 | F: GAGGTGGTAGCAGTAGCAAT |
|  | R: GCAGTGGAGGTTGAGTAGG |
| MSTRG.7888.6 | F: ACAGTCGTCAGTCATCAGTC |
|  | R: TGTGAGTCGTCTATGTTGTCT |
| MSTRG.6027.5 | F: CCAGGCTTGAGCTAACATCT |
|  | R: GATTGAGAGGTGGTGGACAG |
